# Order disorder phase transitions in the plasma membrane of HeLa cells measured by fluorescent analysis of solvatochromic probes

**DOI:** 10.1101/2025.01.29.635609

**Authors:** Nicolas Färber, Sophie C. F. Mauritz, Marina G. Huber, Andrey S. Klymchenko, Christoph Westerhausen

## Abstract

Using the solvatochromic membrane probes Laurdan and Pro12A we investigated order-disorder phase transitions in cellular lipid membranes of HeLa cells. Due to internalization of Laurdan its fluorescence signal yields information about inner and outer cellular membranes while Pro12A exclusively stains the plasma membrane. The two different membrane-embedded dyes show an emission redshift upon increasing disorder of the surrounding lipids that can be quantified by the Generalized Polarization (GP). First, we compare the sensitivity of both probes to lipid phase transitions by analyzing GP of synthetic lipid vesicles as function of temperature. Second, we investigate the temperature dependent lipid order of HeLa cell membranes and find that the plasma membrane shows a partially reversible order-disorder transition regime at temperatures between T = 20 °C and T = 70 °C. Third, we calorimetrically detect an irreversible transition at T = 55 °C and conclude that the optically detected restructuring of the plasma membrane can be partially attributed to protein denaturation. At last, it is shown that the reversible plasma membrane transition temperature T_m_ shifts from T_m_ = 25 °C to T_m_ =-16 °C upon cholesterol depletion and sharpens from a half width of ΔT_FWHM_ = 60 K to ΔT_FWHM_ = 8 K. The reversibility and the sensitivity to cholesterol of this transition indicate a temperature-induced lipid melting within the plasma membrane of HeLa cells.

## 1. Introduction

The lipid structure of cellular membranes is known to have great impact on cellular functions such as membrane transport adhesion, migration, signal transduction and enzyme activity [1–7]. Moreover, it is involved in neuro-degenerative diseases such as Alzheimer’s or Parkinson’s disease [8–10]. Upon distinct diets the lipid composition of colonic caveolae and T-cells can change drastically due to changes in sphingomyelin and cholesterol fractions [11,12]. This lipid structure depends not only on membrane composition but also on environmental parameters such as salt concentration, pH and temperature [13– 15]. All these variables span a multidimensional phase space, in which lipid membranes occur in varying configurations of molecular lipid order. Knowledge on this phase space enables to understand, predict and manipulate the membrane phase state and its associated functions. The lipid structure can be probed using different techniques, such as calorimetry, which works well for simple synthetic lipid systems such as liposomes [13,16]. The analysis of *E. coli* bacterial membranes by Heimburg et al. shows that it can be applied also to more complex biological systems [17]. But our previous studies showed that the temperature dependent lipid order changes in eucaryotic membranes are too small to be detected by calorimetry [13]. This is caused by the high cholesterol content which reduces the cooperativity within lipid membranes and broadens order-disorder phase transitions [18,19]. Other techniques to assess the order state of lipid molecules are Fourier transform infrared spectroscopy, nuclear magnetic resonance spectroscopy and fluorescence spectroscopy [20–22]. The non-linearities of thermodynamic properties of lipid membranes are connected to the sharpness of membrane phase transitions and are key for membrane involved dynamic phenomena, like for instance, the propagation of longitudinal waves in the plasma membrane [23–27]. Recently, it has been demonstrated that order-disorder phase transitions in mammalian cells show broad temperature induced phase transitions for cell ensembles and on the single but whole cell level [1,13]. On the other hand, Fedosejeves et al. reported on sharp phase transitions in living cells from the SH-SY5Y cell line when the analysis is restricted to smaller volumes in the plasma membrane [28]. At first glance, this might be counterintuitive. To address the question, whether there are sharp phase transitions in the plasma membrane of mammalian cells on the level of whole cells, and how sensitive this transition is to the cholesterol level, we chose fluorescence spectroscopy. The latter allows for distinct analysis of different cellular membranes using fluorescent membrane probes with increased affinity for certain membrane regions [29]. The structure of the study is as follows. As probes, we used solvatochromic dyes which indicate an increasing polarity of their environment by a spectral red-shift of their emission spectrum [30]. Solvatochromic dyes, such as Laurdan [31], di-4-ANEPPDHQ [32], Nile Red derivatives [33] etc, have been established as powerful tools to monitor lipid organization, as their emission band position directly correlates with the lipid order [34]. In particular, we have chosen Laurdan and and its derivative Pro12A, which were both specifically designed for the analysis of lipid membrane order [31,35]. Both are incorporated into the bilayer structure and indicate the membrane state following the same principle: Within a highly ordered lipid membrane there is a comparatively small number of solvent molecules present due to high lipid packing density. Upon decreasing lipid order water molecules can penetrate the head group region and the polarity proximal to the fluorescent probes increases. In consequence the emission spectrum shifts to longer wavelengths [36]. The optically determined change of the emission spectrum can be quantified by measuring two wavelength bands and calculating the Generalized Polarization (GP) as shown in **Figure 1** [37]. As Laurdan and Pro12A share the same optically active group they both can be analyzed by determination of GP [35]. Aside from their similar optical properties, Pro12A differs from Laurdan by its membrane anchor composed of dodecyl chain and a negatively charged sulfonate group (**Figure 1**). This membrane anchor ensures that Pro12A exclusively stains the outer leaflet of lipid membranes whereas Laurdan undergoes fast flip-flop and rapid internalization [38,39]. Here, we exploited this property of Laurdan, which is mostly considered a disadvantage and compared lipid order measurements of both probes to distinguish between fluorescent signals from inner and outer membranes of liposomes and cells. The experimental procedure is shown in **Figure 1**. Multilamellar vesicles or HeLa cells were stained by addition of either Pro12A or Laurdan as described in the methods sections **2.3** and **2.4**. As a result, either all membranes of the different lipid systems were stained or only the outer membranes. Afterwards, the bulk sample suspension was analyzed in a reaction tube at different temperatures using a ultraviolet LED for excitation and a spectrometer to record the spectra. Using this approach, we obtained GP temperature profiles which allow for determination of the lipid system phase state and to detect phase transitions between different degrees of molecular order.

**Figure 1.**
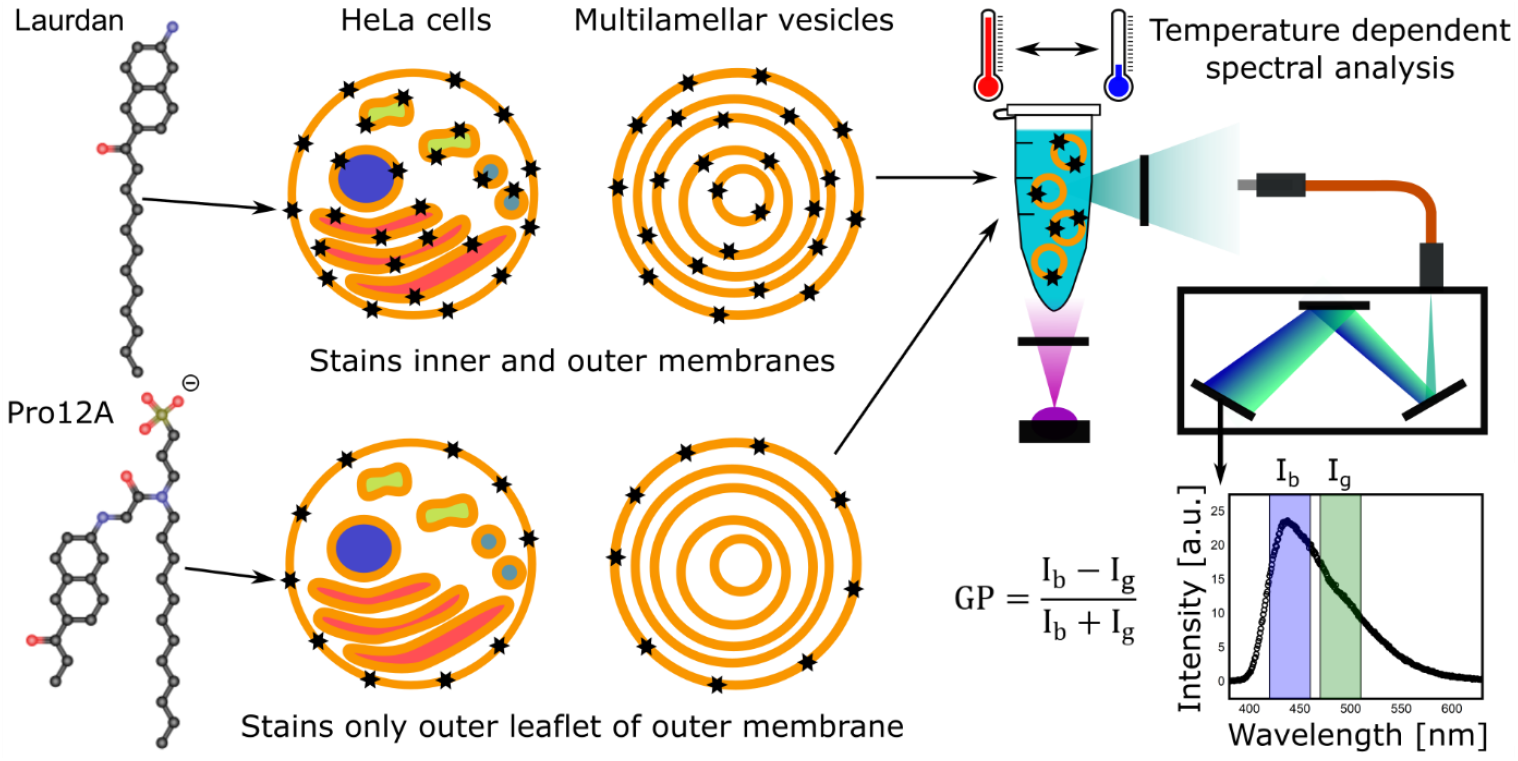
Schematic representation of the experimental design. Two membrane-embedded probes were used to analyze the lipid order state of HeLa cell membranes and multilamellar vesicles. Upon addition to the suspended vesicles or cells Laurdan stains inner and outer lipid membranes due to internalization, while Pro12A labels only the outer leaflet of the outer membrane because of its negatively charged sulfonate membrane anchor. After staining the sample suspensions were analyzed in reaction tubes at different temperatures using an ultraviolet (UV) LED with UV-bandpass filter for dye excitation and a spectrometer with an UV-band stop filter to collect the emitted light. The intensities of two wavelength regions centred around λ = 440 nm and λ = 490 nm were integrated to calculate the Generalized Polarization (GP) as measure for the mean lipid order of a sample suspension.

## 2. Materials and methods

In this study we investigated synthetic and biological lipid systems using fluorescence spectroscopy and calorimetry. The preparation of these systems, the measurement techniques and the data analysis are described in the following passage.

### 2.1 Preparation of multilamellar vesicles

2 mM of either 14:0 PC or 15:0 PC (Sigma-Aldrich Chemie GmbH, Munich, Germany) phospholipids dissolved in chloroform were dried under nitrogen flow in 4 ml glass vials followed by vacuum exposure for 60 min. Vesicle swelling was induced by addition of 1 ml ultra-pure water and ultrasonic treatment for 30 min at a temperature of 40 °C (14:0 PC) or 50 °C (15:0 PC). The final lipid concentration was 2 mM.

### 2.2 Culture of HeLa cells

HeLa cells (ATCC® CCL-2™) were cultured in 25 cm^2^ Nunc™ cell culture flasks (ThermoFisher Scientific, MA, USA) at 37 °C in saturated atmosphere in DMEM (Bio&SELL GmbH, Nürnberg, Germany) supplemented with 10 % fetal bovine serum (FBS Superior) and 1 % penicillin / streptomycin (Biochrom GmbH, Berlin, Germany). 5 % of cells were passaged using 0.05 % trypsin/EDTA (Biochrom GmbH) each time when a confluency of 80 % was reached.

### 2.3 Staining of multilamellar vesicles

Laurdan (Sigma-Aldrich Chemie GmbH, Munich, Germany) and Pro12A (Laboratoire de Biophotonique et Pharmacologie of Andrey Klymchenko, Université de Strasbourg, France) [35] were dissolved in dimethyl sulfoxide (DMSO) at a dye concentration of 20 µM. The staining solution was added in a volume ratio of 1:50 to the vesicle suspensions resulting in a final dye concentration of 0.4 µM and a final DMSO concentration of 276 mM (21.6 mg/ml). Laurdan was added to the vesicle suspension at least 120 min prior to spectroscopic analysis to ensure that the internalization process is completed. This equilibration time was not necessary in case of staining with Pro12A as it is not internalized. We did not identify a change of the fluorescent signal after addition on experimental time scales of a few minutes.

### 2.4 Staining of HeLa cell membranes

1 mg of Laurdan was dissolved in 1 ml of DMSO yielding a dye concentration of 2.8 mM. This stock solution was added in a volume ratio of 1:50 into the culture medium of the adherent HeLa cells yielding a final concentration of 56 µM Laurdan and 276 mM DMSO. After two hours of incubation at 37 °C the medium was removed, and the cells were detached using 1ml of 0.05 % trypsin/EDTA solution. The as obtained cell suspension was immediately analyzed. In case of Pro12A staining there was no incubation procedure necessary. After detachment using 1ml of 0.05 % trypsin/EDTA solution the Pro12A stock solution (0.5 mM in DMSO) was added to the suspended cells in a volume ratio of 1:50. The final concentrations were 10 µM Pro12A and 276 mM DMSO. After dye addition the cell suspension was immediately analyzed. Each sample suspension contained about 2 Mio cells.

### 2.5 Cholesterol depletion of HeLa cell membranes

The cholesterol content of cell membranes was reduced by exposure to Methyl-β-cyclodextrin (MBCD). First, adherent cells were detached with 1ml of 0.05 % trypsin/EDTA solution and the resulting suspension was centrifuged at 300 G for 5 min. Afterwards the supernatant was removed and the cell pellet was resuspended in 0.6 ml phosphate buffered saline (PBS) containing 10 mM MBCD. Pro12A was added to this cell suspension as described above followed by immediate analysis.

### 2.6 Calorimetric analysis of cell and vesicle suspensions

Differential scanning calorimetry (DSC) was performed using a MicroCal VP-DSC calorimeter (MicroCal Inc., now Malvern Panalytical Ltd.,UK). Multilamellar vesicles were analyzed in degassed ultrapure water at 15 psi pressure with a scan rate of 10 K/h and ultrapure water as reference. The down-scan after three subsequent measurement cycles was used for data analysis and corrected by baseline subtraction as described previously [40]. For the analysis of biological membranes we cultured HeLa cells in 175 cm^2^ Nunc™ cell culture flasks (ThermoFisher Scientific, MA, USA). The cells of two fully grown flasks were collected using 0.05 % trypsin/EDTA solution followed by centrifugation. After supernatant removal the pellet (about 28 Mio. cells) was resuspended in 1 ml PBS solution and measured against a PBS solution reference.

### 2.7 Spectroscopic analysis of cell and vesicle suspensions

The stained cell suspensions were transferred into a 600 µl reaction tube, that was fitted into an aluminum block. This block was tempered using a combination of a thermoelectric element and a water cooling system. The temperature was recorded *in-situ* with an encapsulated Pt100-sensor that was inserted through the lid of the reaction tube. The optical properties were assessed through boreholes in the aluminum block. We used a 360 nm LED (Power 1 W)in combination with a 360 nm bandpass filter <u>(Δλ = 30 nm)</u> to excite the membrane-embedded probes. After passing a 360 nm band stop filter the fluorescent light was collected using a 1 mm diameter optical fiber attached to an ocean optics QEPro spectrometer. Measurements were taken in a range from T =-35 °C to T = 90 °C using a heating/cooling rate of 2 K/min. Spectra were recorded in steps of ΔT = 1 K. The whole setup was controlled using a LabView program that enabled fully automatic temperature scans.

### 2.8 Analysis of spectral data

For every spectrum we calculated the GP value according to the equation

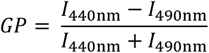

(**Figure 1**) using the integrated intensities I_blue_ and I_green_ over a wavelength region of λ = 420 nm to λ = 460 nm and λ = 470 nm to λ = 510 nm. For one temperature point three GP values of independently conducted experiments were averaged and the standard deviation was calculated as an error indicator.

## 3. Results and Discussion

The study is structured as follows: We start with the calorimetric and fluorescent analysis of synthetic model membranes to identify differences in the optical properties of Pro12A and Laurdan (**3.1**). Afterwards we apply a similar data analysis to optical data of HeLa cell membranes and identify two distinct transition regimes (**3.2**) that we investigate further in the last two sections. First, we apply differential scanning calorimetry (**3.3**). Second, we reduce the cholesterol within the cellular membranes (**3.4**).

### 3.1. Order-disorder transitions in synthetic lipid membranes

Prior to the evaluation of the complex lipid system of biological membranes, we compared the optical properties of the two membrane probes Pro12A and Laurdan by analyzing well-defined model systems. We chose multilamellar vesicles as they have inner and outer membranes, similar as cells, that can be targeted as a whole for Laurdan or exclusively the outer membranes using Pro12A. The multilamellar vesicles were obtained by hydration of lipid films (see 2.1). After the preparation, the liposomes were stained by addition of the DMSO dissolved membrane probes, as described in 2.3. The emission spectra at T = 5 °C and T = 55 °C of Laurdan and Pro12A embedded in 14:0 PC multilamellar vesicles are depicted in **Figure 2 a)**. These vesicles exhibit a transition from the ordered gel-like phase to the disordered fluid phase at T_m_ = 24 °C, as can be seen in the calorimetric data presented in **Figure 2 b)**. Therefore, the data show the spectral response of the membrane probes to a lipid membrane phase state change. The maximum fluorescence intensity of membrane-embedded Laurdan is located at a wavelength of λ = 449 nm corresponding to a fully ordered membrane. This value is slightly higher than the literature value of λ = 440 nm which could be caused by a spectral offset of the used spectrometer [37]. Upon heating and undergoing a transition to the fluid membrane phase, the emission spectrum is red shifted and the maximum fluorescence intensity at T = 55 °C is located at λ = 495 nm. Again this value is higher than the literature value of λ = 490 nm observed by Parasassi et al. [37]. In consequence, we observed a maximum Stokes-shift of Laurdan induced by a lipid melting transition of Δλ = 46 nm. Even though Laurdan and Pro12A have the same optically active naphthalene group the emission spectra of the two probes differ within the same lipid system. In **Figure 2 a)** the highest emission intensity of Pro12A at T = 5 °C is observed at λ = 465 nm. The different fluorescent signals might be explained by different locations of the membrane probes within the bilayer which is known to influence the emission properties of solvatochromic dyes [41]. The anionic anchor of Pro12A could cause a location closer to the lipid head group water interface where the membrane hydration level is higher. As a result, the fluorescence emission of Pro12A would be red-shifted compared to the less hydrated Laurdan, due to increased solvent relaxation. An increased hydration level within the local environment of Pro12A might also be caused by induction of a larger membrane defect because of the size difference of the two membrane probes. However, in the fluid phase at T = 55 °C the emission maximum of Pro12A is less red-shifted compared to Laurdan and can be observed at λ = 478 nm. This indicates that the optical active area of Pro12A senses a less hydrated environment in a fluid membrane as compared to Laurdan. The higher spectral response of Laurdan could be explained by a relocation of the dye towards the surface induced by the structural phase transition of the surrounding lipid system. Such a relocation caused by conformational changes of Laurdan were recently observed by Bacalum et al. [42]. Pro12A bearing the anchor group cannot really undergo this relocation, so its spectral response is smaller. The maximal observed Stokes-shift of Pro12A is Δλ = 13 nm. Therefore, it shows less spectral response to melting transitions within multilamellar vesicles as compared to Laurdan. To investigate in detail the capability of optical phase transition analysis by the fluorescent membrane probes, we recorded emission spectra at various temperatures above and below the melting transition of two different vesicle samples. Their temperature induced phase state change is indicated by the maxima in the heat capacity profile that is shown in **Figure 2 b)**. To quantitate the optical determined phase state, we calculated the GP value for each spectrum. The corresponding values are depicted in **Figure 2 c)** as function of temperature for both lipid systems measured using the membrane probe Laurdan. Below the melting points of T_m_ = 24 °C (14:0 PC) and T_m_ = 34 °C (15:0 PC) the GP values of both lipid systems are positive and 15:0 PC vesicles show a slightly higher GP level. At the respective transition temperature, a sharp drop of GP can be observed followed by a steady decrease towards values of GP = 0.4. The negative derivatives with respect to temperature of the GP graphs also shown in **Figure 2 c)** indicate the phase transitions as a sharp maximum similar to the calorimetric measurements depicted in **Figure 2 b)**. We have shown the correlation of the quantity-dGP/dT and the heat capacity profile of lipid systems before [13,16]. The temperature dependent fluorescent analysis of solvatochromic dyes can be used as complementary measurement to differential scanning calorimetry. The equivalent data analysis of Pro12A is presented in **Figure 2 d)**. In contrast to the Laurdan data, the GP values of Pro12A embedded in 15:0 PC are lower than for 14:0 PC. Furthermore, a less steep transition within the gel-phase can be observed around temperatures of T = 9 °C and T = 18 °C. These could be attributed to the pre-transition from the gel-like phase into the ripple that are also visible in the calorimetric data shown in **Figure 2 b)**. At the main melting transition both samples exhibit a drop of GP, as observed for Laurdan. The transition is not as sharp as in **Figure 2 b)** causing the-dGP/dT values to be smaller as compared to Laurdan. In agreement with the reduced Stokes-shit of Pro12A shown in **Figure 2 a)** the GP range covered in the Temperature interval from T = 5 °C to T = 55 °C is smaller (ΔGP ∼ 0,45) as compared to Laurdan (ΔGP ∼ 0,65). In summary Pro12A, is less sensitive to the phase state of multilamellar vesicles as compared to Laurdan but both probes allow for clear detection of lipid membrane melting.

**Figure 2.**
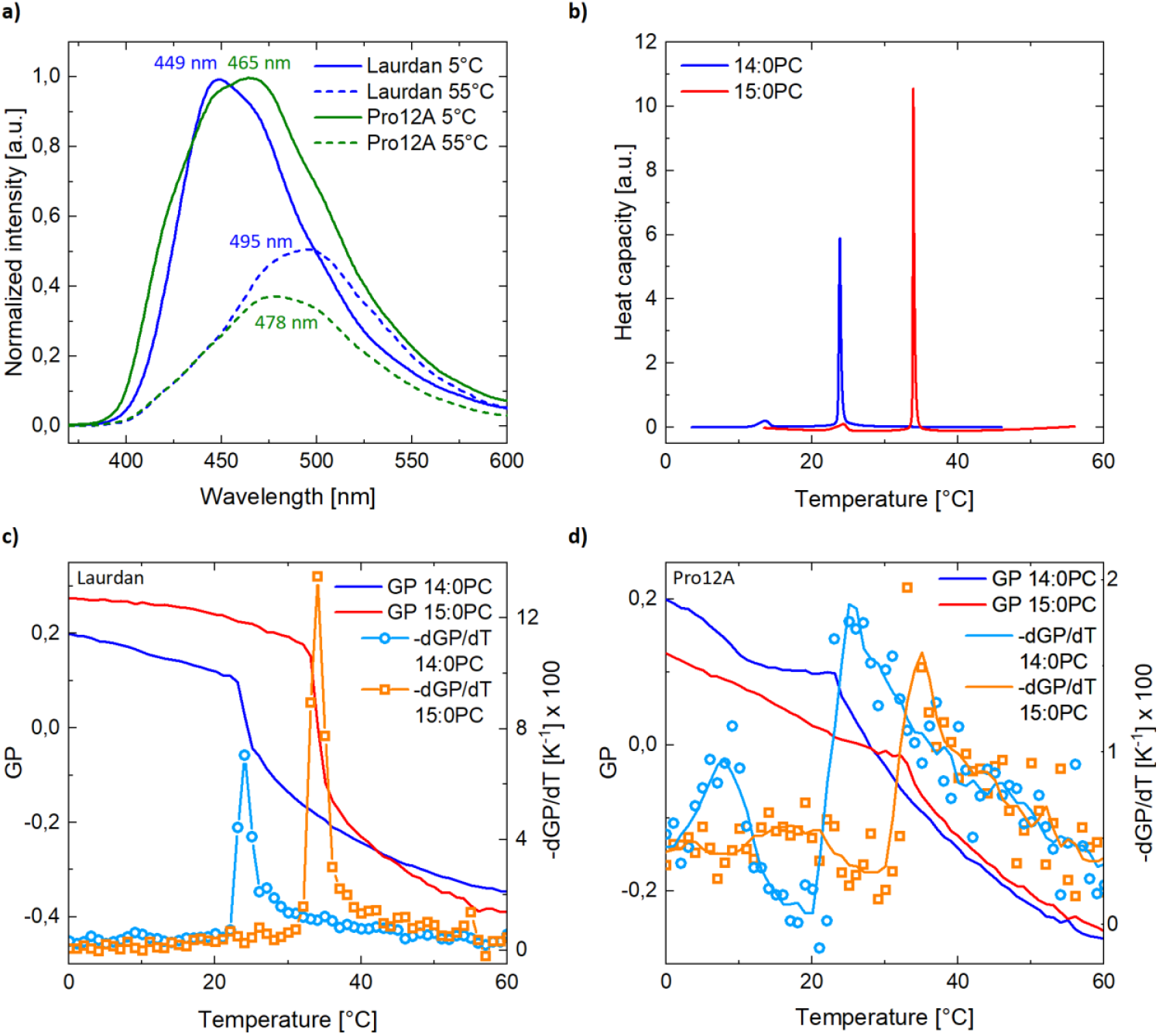
Optical lipid order and calorimetric measurements on multilamellar lipid vesicles. **a)** Normalized emission spectra of Laurdan and Pro12A embedded in lipid membranes of multilamellar 14:0 PC vesicles at T = 5 °C and T = 55 °C. The spectral red shift upon increased temperature indicates increased lipid disorder. **b)** Downscan heat capacity of multilamellar 14:0 PC and 15:0 PC vesicles. The maxima indicate lipid membrane phase transitions from the disordered or fluid phase to the ordered or gel-like phase. **c)** The lipid order of multilamellar 14:0 PC and 15:0 PC vesicles measured by fluorescent analysis of membrane-embedded Laurdan and quantified by GP as function of temperature during downscanning. As Laurdan is internalized into inner membranes of the multilamellar vesicles the resulting fluorescence signal originates from all vesicle membranes. The sharp decrease of GP indicates the phase transitions that were calorimetrically measured in b). The negative derivatives of GP with respect to temperature show pronounced maxima at the transition from the ordered to the disordered phase. **d)** GP and its negative derivative as function of temperature as presented in c) but measured using membrane-embedded Pro12A instead of Laurdan. This membrane probe stains exclusively the outer leaflet of the outer vesicle membranes in contrast to Laurdan. A decrease of GP indicates the phase transitions as seen for Laurdan but the step-like behaviour is not as pronounced. In consequence the maxima of the negative derivative of GP are also not that sharp as in c). Therefore-dGP/dT (single dots) was smoothed using a flying average (solid line).

### 3.2 Order-disorder transitions in HeLa cell membranes

Analogously to the described analysis of multilamellar vesicles we recorded spectra for membrane-stained HeLa cells as function of temperature. Prior to this analysis the cells were cultured and stained as described in the methods section **2.2** and **2.4**. GP as function of temperature during heating the cell membranes from T = 5 °C to T = 90 °C is shown in **Figure 3 a)** for Pro12A and Laurdan stained membranes. Even though such high temperatures do not have a physiological equivalent, we chose this wide temperature range to cover the full transition regime of the lipid membrane system. The comparison of both measurements yields a higher GP value for Pro12A stained cells as compared to Laurdan-labeled samples especially at lower temperatures. This is in agreement with prior findings that inner cell membranes of organelles are more fluid than the plasma membrane [18,43,44]. As the fluorescence signal of Laurdan originates from both outer and inner membranes, the GP value is in consequence lower than for Pro12A that probes only the outer leaflet of the plasma membrane [35]. Furthermore, the GP-profiles shown in **Figure 3 a)** differ in their temperature dependent GP change. Especially around T = 55 °C the GP profile of Pro12A stained HeLa cell membranes is steeper as compared to Laurdan. This is also reflected in the derivative with respect to temperature shown in **Figure 3 c)**. In contrast to the derivative-dGP/dT of Laurdan that does not display a distinct transition regime the Pro12A stained membranes exhibit a pronounced maximum at T = 55 °C. Moreover, a second regime can be observed at lower temperatures that extends into the other maximum. The investigation of the origin of these two transitions will be part of section **3.3** and **3.4**. The heating of the samples was followed by cooling from T = 90 °C to T =-30 °C. The corresponding GP profiles during the down-scan are shown in **Figure 3 b)**. The GP profile of Laurdan stained HeLa cell membranes is shifted over the whole temperature regime towards higher GP values indicating an increased lipid order. For example, GP ∼-0.2 during up-scanning at T = 30 °C and GP ∼ 0 in the down-scan. This might be explained by a conformational change of the membrane proteins upon heat induced denaturation. It was shown using molecular dynamics simulations that the order state of lipids is affected by the conformation of proximal proteins [45]. A similar effect can be observed for Pro12A marked HeLa cell membranes: The membrane order decreases from GP ∼ 0.2 during the up-scan to GP ∼ 0 for the denaturated membranes in the down-scan. Both membrane-probes indicate a denaturation-induced change of lipid order. The inner membrane compartments probed by Laurdan exhibit increased lipid order while the order of the plasma membrane probed by Pro12A is decreased. In addition to protein denaturation the reduction of GP in case of Pro12A after heating could be partially caused by redistribution of the membrane probe. It can not be excluded that Pro12A is internalized at elevated temperatures resulting in fluorescent signal from inner cell membranes with lower GP-value. The temperature dependence of lipid order during the down-scan is shown in **Figure 3 d)**. In case of Laurdan the - dGP/dT graph displays a clear lipid order transition with a maximum around T = 15 °C. This transition could not be observed in **Figure 3 c)**, which could be caused by a shift of the melting transition towards higher temperatures due to heat induced protein denaturation. The melting transition in Pro12A membranes located at T = 55 °C **Figure 3 c)** disappeared after heating to T = 90 °C as can be seen in **Figure 3 d)**. Only a less cooperative transition can be observed at T = 30 °C which could be attributed to the left shoulder visible in the double peak structure in **Figure 3 c)**. The behaviour of the lipid order transition within the plasma membranes of HeLa cells could be explained by a reversible phase transition of the membrane lipid system. The irreversible part might be attributed to a phase state change due to a change of membrane protein conformation induced by heat denaturation. To further investigate the origin of the observed plasma membrane phase transitions we measured the heat capacity of whole HeLa cells. Prior experiments showed that lipid phase transitions within eukaryotic cell membranes can hardly be detected by differential scanning calorimetry due to their high cholesterol content and low cooperativity. But if the lipid order transitions should be induced by membrane protein denaturation, the corresponding protein melting event should be detectable during the first calorimetric up-scan.

**Figure 3.**
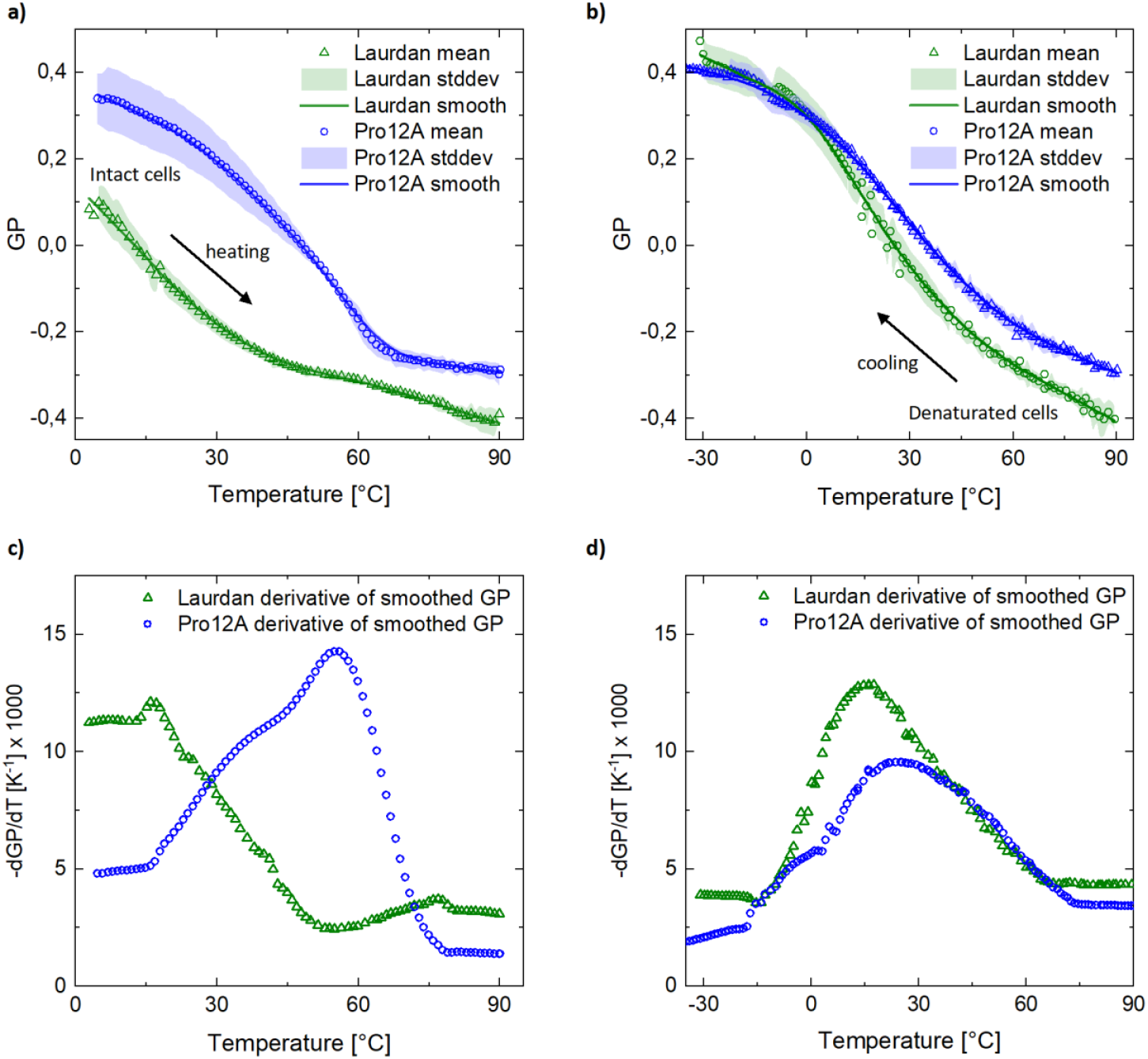
Optical lipid order measurements on HeLa cell membranes using Laurdan and Pro12A. While Laurdan stains inner and outer cellular membranes due to internalization the fluorescent signal of Pro12A originates primarily from the outer leaflet of the plasma membrane. **a)** The lipid order quantified by GP of HeLa cell membranes measured by fluorescent analysis of Laurdan and Pro12A during heating from T = 5 °C to T = 95 °C. The data points represent the mean values of three independently conducted experiments. While GP measured using Laurdan shows a continuous decrease with respect to temperature the Pro12A graph exhibits a stronger drop at T = 60°C. Furthermore, the absolute GP values of Pro12A stained cellular membranes are higher as compared to Laurdan. This indicates that the plasma membrane of Hela cells is in a phase state of higher molecular order and it exhibits a higher temperature sensitivity as compared to inner cellular membranes. **b)** GP measured from T = 90 °C to T =-30 °C of the cellular membranes analysed in a). The data points represent the mean values of three independently conducted experiments. In contrast to the upscans shown in a) the fluorescent signal originates from membranes that are denaturated due to high temperatures. The comparison of the lipid order in the plasma membrane measured by Pro12A and the signal of Laurdan representing a mean value of inner and outer membranes shows less difference as the data of intact membranes shown in a). Both GP profiles are located at higher GP values but the increase of lipid order in the plasma membrane is much higher than the overall lipid order measured by Laurdan. Furthermore, the GP temperature dependence of Laurdan-stained HeLa cell membranes is higher as compared to the Pro12A samples. **c)** The negative derivative with respect to temperature of the loess-smoothed GP values shown in a). While the sum of inner and outer cellular membranes measured by Laurdan exhibits no distinct maximum the Pro12A analysis of the plasma membrane shows a broad transition regime that extends from T = 20 °C to T = 70 °C. Furthermore, two shoulders are visible that could originate from two different processes of lipid order rearrangement. **d)** The negative derivative with respect to temperature of the loess-smoothed GP values shown in b). The right shoulder of the plasma membrane transition shown in c) disappears after denaturation at high temperatures indicating an irreversible process of membrane reordering. The left shoulder of the transition can be attributed to a reversible process. While no distinct transition could be seen in c) using Laurdan there is an even more pronounced transition after denaturation as compared to the plasma membrane.

### 3.3 Calorimetric analysis of transitions in HeLa cells

Similar to the optical scan shown in **Figure 3 c)** we analyzed the heat capacity of whole HeLa cells during up-scanning as can be seen in **Figure 4 a)**. Three samples were measured, and the baseline was approximated by a cubic function. The baseline corrected heat capacity profiles are shown **Figure 4 b)**. All samples exhibit a maximum heat capacity slightly below T = 60 °C. As this transition was not observed in the following scans the origin of the calorimetrically detected transition must by an irreversible process. These findings are in agreement with previous calorimetric studies on whole HeLa cells which also exhibited protein-associated melting transitions near T = 60 °C [46]. The comparison of the optically detected phase transition using Pro12A, and the summarized calorimetric analysis is shown in **Figure 4 c)**. This data suggests that the irreversible lipid order transition within the plasma membrane measured using Pro12A could be caused by the denaturation of membrane associated protein. The left shoulder of the optically detected transition that is also visible in the down-scan as shown in **Figure 4 d)** could indicate a reversible lipid melting within the plasma membrane. If this was the case, the reversible transition should be sensitive to the membrane composition. For instance it is known that cholesterol strongly influences the phase state of synthetic and biological membranes [47]. In the following section we investigated whether the manipulation of the HeLa cell membrane cholesterol content would affect the optically observed reversible plasma membrane transition shown in **Figure 4 d)**.

**Figure 4.**
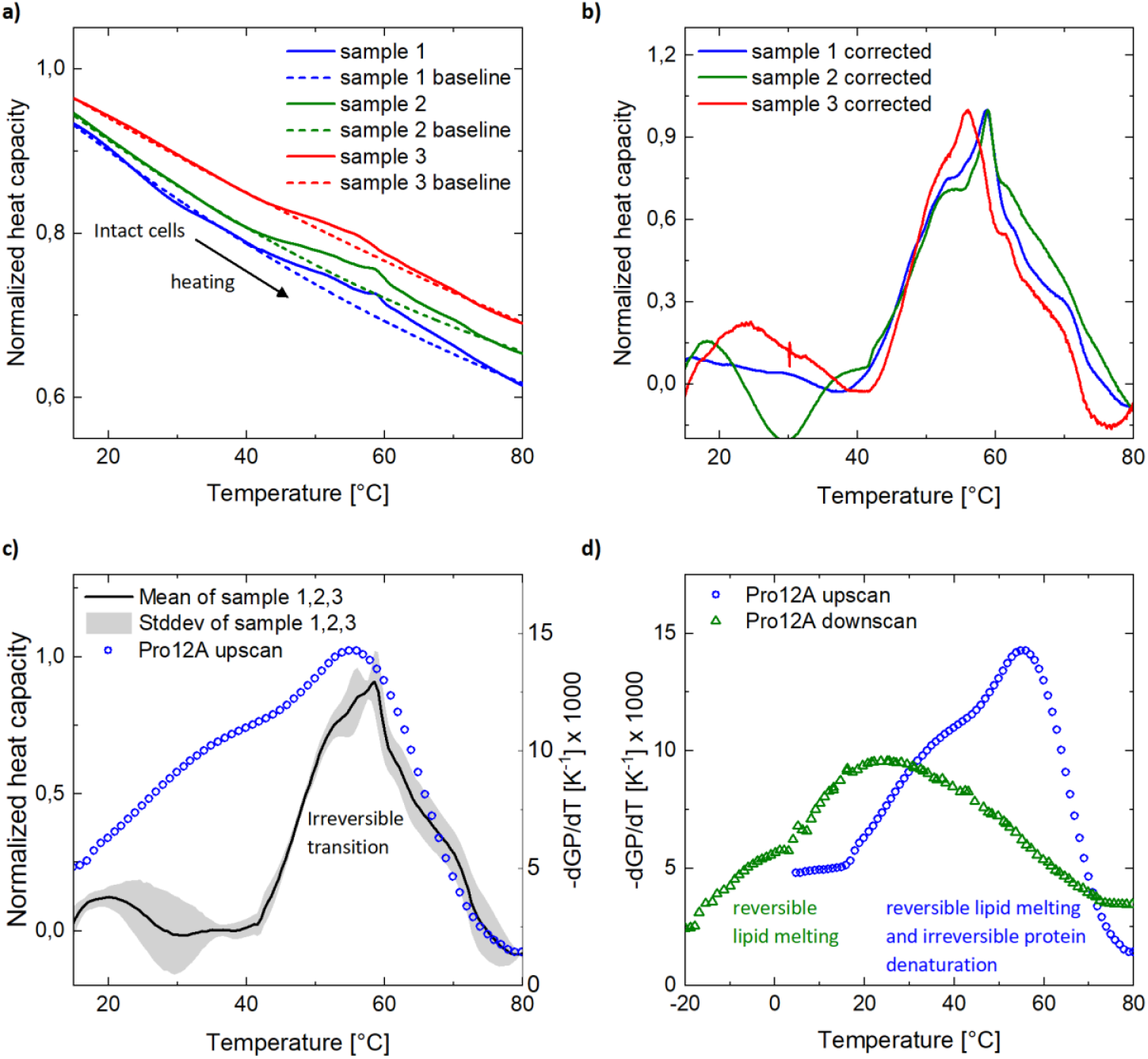
Calorimetric analysis of whole HeLa cells and comparison to optical plasma membrane data obtained using Pro12A. **a)** Heat capacity profiles (solid lines) of three independently prepared HeLa cell suspensions measured from T = 15 °C to T = 80 °C. Each dataset was normalized to its maximum value and the baselines were approximated by cubic functions (dotted lines). **b)** The normalized heat capacity profiles of a) corrected for the corresponding baselines. All samples show an irreversible melting event around T = 55 °C that could not be observed in the subsequently conducted scans. **c)** Comparison of the mean normalized heat capacity profiles of whole HeLa cells obtained at first upscan and the lipid plasma membrane phase transition of HeLa cells measured in the first upscan using Pro12A. Both calorimetric and optical data show an irreversible transition around T = 55 °C. This indicates that the right shoulder of the optically measured plasma membrane transition could be explained by a membrane restructuring due to membrane protein denaturation. **d)** Comparison of optically determined plasma membrane transitions measured during heating and cooling of cellular membranes. The reversible transition regime that is left after denaturation could be attributed to lipid melting as part of a phase transition within the plasma membrane.

### 3.4 Plasma membrane sensitivity to cholesterol depletion

To exclusively analyze the reversible lipid order transition of HeLa plasma membranes, Pro12A spectra were recorded during cooling from T = 90 °C to T =-35 °C. The influence of cholesterol content on the phase transition was investigated using MBCD as described in **2.5**. The comparison of MBCD treated membranes with reduced cholesterol level and an untreated control are shown in **Figure 5 a)**. The GP value is strongly decreased over the whole measured temperature range corresponding to a reduction of lipid order. This is in agreement with previous studies showing that cholesterol has an ordering effect within lipid membranes [48]. This effect is much more pronounced than the GP changes upon cholesterol extraction that we investigated in an earlier study using Laurdan and thus probing all membrane compartments of the cell [13,35]. This can be explained by the larger cholesterol content within the plasma membrane as compared to inner cellular membranes [18]. Most interestingly the MBCD treated sample in **Figure 5 a)** exhibits a kink in the GP curve at T =-16 °C indicating a sudden change of lipid order. This results in a sharp maximum of the derivative of GP with respect to temperature shown in **Figure 5 b)**. The comparison of both graphs yields that the reversible phase transition within plasma membranes of HeLa cells shifts from a temperature of T = 25 °C to T =-16 °C. Furthermore, it sharpens from a transition half width of ΔT_FWHM_ = 60 K to ΔT_FWHM_ = 8 K upon cholesterol extraction. Both results are in agreement with previous studies with lipid vesicles showing a broadening and a shift of the melting temperature with increasing levels of cholesterol [49,50]. These findings highlight the importance of cholesterol for the plasma membrane phase state. It ensures that the bilayer melting temperature is located right below the culture temperature under physiological conditions. Previous studies have shown the proximity of membranes to phase transitions for various organisms before [17,51,52]. Variation of the cholesterol fraction in the plasma membrane changes the non-linear mechanical properties of the membrane dramatically. Our findings suggest that it plays a key role to sustain the phase state dependent physical properties of the membrane.

**Figure 5.**
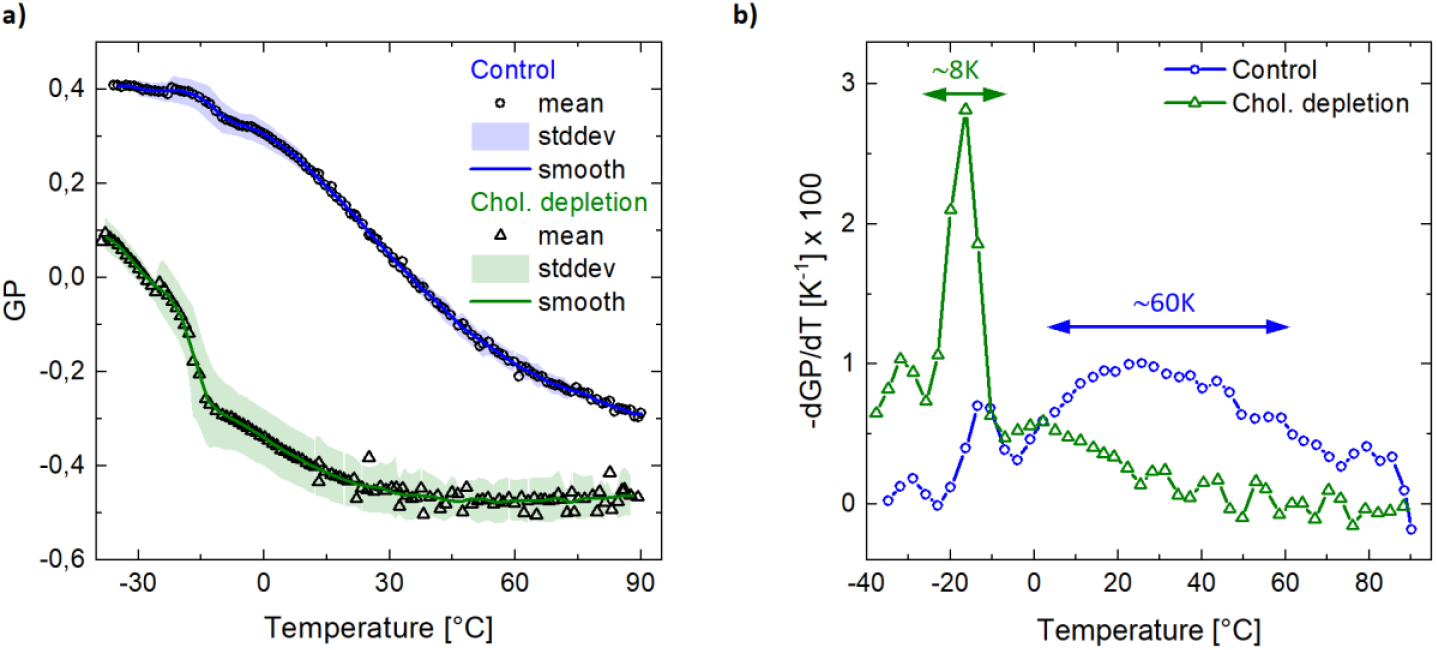
Lipid plasma membrane order of HeLa cells measured using Pro12A before and after cholesterol depletion. **a)** GP as measure for the lipid plasma membrane order recorded during cooling from T = 90 °C to T =-35 °C. The data points represent the mean value of three independently conducted experiments. The plasma membrane order is strongly decreased over the whole temperature regime upon cholesterol extraction. Furthermore, the temperature dependence of the lipid order is increased at temperatures below zero as compared to the control measurement. **b)** The comparison of the negative derivatives of GP with respect to temperature yields a narrowing of the transition regime from ΔT_FWHM_ = 60 K to ΔT_FWHM_ = 8 K and a shift of the melting temperature T_m_ from Tm = 25 °C to Tm =-16 °C upon cholesterol depletion.

## 4. Conclusion

We demonstrated for multilamellar vesicles that the temperature dependent fluorescent analysis of Pro12A and Laurdan yields the lipid order state and membrane melting transitions similar to calorimetry. Furthermore, the same concept can be applied to cellular membranes where the comparison of Laurdan and Pro12A allows to differentiate between lipid phase transitions within the plasma membrane and the membranes of inner cell organelles. We found that in contrast to the whole cell analysis, Pro12A resolves two distinct lipid order transitions within the plasma membrane. One of them is irreversible and is most likely caused by protein denaturation. The other is reversible and exhibits a high sensitivity to the plasma membrane cholesterol content. Upon cholesterol depletion, this transition shifts into the negative temperature regime and sharpens to a transition half width of ΔT_FWHM_ = 8 K. Our findings suggest that cholesterol fulfils the role of adjusting the melting regime of the plasma membrane in a way that the overall cell is living on the fluid side of a broad phase transition.

## Acknowledgements

N.F. thanks the Joachim Herz Stiftung for financial support. C.W. would like to acknowledge funding by the DFG (INST 94/135-1 FUGG, 507881424 and 508235635). Moreover, N.F. and C.W. thank the Augsburg Centre of Innovative Technologies (ACIT) for support and Achim Wixforth for fruitful discussions.

## Data availability

All data that support our findings are available from the corresponding authors upon reasonable request. There are no restrictions on data availability.

## Author contributions

N.F., A.S.K, and C.W. designed research; N.F., S.M., M. H. and C.W. performed research; N.F., S.M. and M.H. analyzed data; N.F. implemented experimental tools; N.F. and C.W. wrote the original manuscript. A.S.K. provided the fluorescent probe and discussed the results. All authors worked on the final manuscript.

## References

[1] Färber N, Reitler J, Schäfer J, Westerhausen C. Transport Across Cell Membranes is Modulated by Lipid Order. Adv Biol 2023:2200282.

[2] Ikonen E. Roles of lipid rafts in membrane transport. Curr Opin Cell Biol 2001;13:470–7. 10.1016/S0955-0674(00)00238-6.

[3] Gaus K, Le Lay S, Balasubramanian N, Schwartz MA. Integrin-mediated adhesion regulates membrane order. J Cell Biol 2006;174:725–34. 10.1083/jcb.200603034.

[4] Braig S, Sebastian Schmidt BU, Stoiber K, Händel C, Möhn T, Werz O, et al. Pharmacological targeting of membrane rigidity: implications on cancer cell migration and invasion. New J Phys 2015;17:83007. 10.1088/1367-2630/17/8/083007.

[5] Fichtl B, Silman I, Schneider MF. On the Physical Basis of Biological Signaling by Interface Pulses. Langmuir 2018;34:4914–9. 10.1021/acs.langmuir.7b01613.

[6] Fichtl B, Shrivastava S, Schneider MF. Protons at the speed of sound: Predicting specific biological signaling from physics. Sci Rep 2016;6:1–9. 10.1038/srep22874.

[7] Kamenac A, Obser T, Wixforth A, Schneider MF, Westerhausen C. The activity of the intrinsically water-soluble enzyme ADAMTS13 correlates with the membrane state when bound to a phospholipid bilayer. Sci Rep 2021;11:24476.

[8] Fabelo N, Martín V, Santpere G, Marín R, Torrent L, Ferrer I, et al. Severe alterations in lipid composition of frontal cortex lipid rafts from Parkinson’s disease and incidental Parkinson’s disease. Mol Med 2011;17:1107–18.

[9] Gasecka P, Jaouen A, Bioud F-Z, de Aguiar HB, Duboisset J, Ferrand P, et al. Lipid order degradation in autoimmune demyelination probed by polarized coherent Raman microscopy. Biophys J 2017;113:1520–30.

[10] Martín V, Fabelo N, Santpere G, Puig B, Marín R, Ferrer I, et al. Lipid Alterations in Lipid Rafts from Alzheimer’s Disease Human Brain Cortex. J Alzheimer’s Dis 2010;19:489–502. 10.3233/JAD-2010-1242.

[11] Ma DWL, Seo J, Davidson LA, Callaway ES, Fan Y-Y, Lupton JR, et al. n-3 PUFA alter caveolae lipid composition and resident protein localization in mouse colon. FASEB J 2004;18:1040–2.

[12] Fan Y-Y, McMurray DN, Ly LH, Chapkin RS. Dietary (n-3) polyunsaturated fatty acids remodel mouse T-cell lipid rafts. J Nutr 2003;133:1913–20.

[13] Färber N, Westerhausen C. Broad lipid phase transitions in mammalian cell membranes measured by Laurdan fluorescence spectroscopy. Biochim Biophys Acta - Biomembr 2022;1864:183794. 10.1016/j.bbamem.2021.183794.

[14] Busto J V., García-Arribas AB, Sot J, Torrecillas A, Gómez-Fernández JC, Goñi FM, et al. Lamellar gel (Lβ) phases of ternary lipid composition containing ceramide and cholesterol. Biophys J 2014;106:621–30. 10.1016/j.bpj.2013.12.021.

[15] Trauble H, Eibl H. Electrostatic effects on lipid phase transitions: membrane structure and ionic environment. Proc Natl Acad Sci U S A 1974;71:214–9. 10.1073/pnas.71.1.214.

[16] Färber N, Reitler J, Kamenac A, Westerhausen C. Shear stress induced lipid order and permeability changes of giant unilamellar vesicles. Biochim Biophys Acta (BBA)-General Subj 2022;1866:130199.

[17] Mužić T, Tounsi F, Madsen SB, Pollakowski D, Konrad M, Heimburg T. Melting transitions in biomembranes. Biochim Biophys Acta - Biomembr 2019;1861:183026. 10.1016/j.bbamem.2019.07.014.

[18] Yeagle PL. Cholesterol and the cell membrane. Biochim Biophys Acta (BBA)-Reviews Biomembr 1985;822:267–87.

[19] Redondo-Morata L, Giannotti MI, Sanz F. Influence of cholesterol on the phase transition of lipid bilayers: a temperature-controlled force spectroscopy study. Langmuir 2012;28:12851–60.

[20] Lewis RNAH, McElhaney RN. Membrane lipid phase transitions and phase organization studied by Fourier transform infrared spectroscopy. Biochim Biophys Acta - Biomembr 2013;1828:2347–58. 10.1016/j.bbamem.2012.10.018.

[21] Leung SSW, Brewer J, Bagatolli LA, Thewalt JL. Measuring molecular order for lipid membrane phase studies: Linear relationship between Laurdan generalized polarization and deuterium NMR order parameter. Biochim Biophys Acta - Biomembr 2019;1861:183053. 10.1016/j.bbamem.2019.183053.

[22] Niko Y, Klymchenko AS. Emerging solvatochromic push–pull dyes for monitoring the lipid order of biomembranes in live cells. J Biochem 2021;170:163–74.

[23] Fox D. The Brain, Reimagined. Sci Am 2018;318:60–7. 10.1038/scientificamerican0418-60.

[24] Heimburg T, Jackson AD. On soliton propagation in biomembranes and nerves. Proc Natl Acad Sci U S A 2005;102:9790–5. 10.1073/pnas.0503823102.

[25] Lautrup B, Appali R, Jackson AD, Heimburg T. The stability of solitons in biomembranes and nerves. Eur Phys J E 2011;34:1–9.

[26] Heimburg T. The thermodynamic soliton theory of the nervous impulse and possible medical implications. Prog Biophys Mol Biol 2022;173:24–35.

[27] Mussel M, Schneider MF. Sound pulses in lipid membranes and their potential function in biology. Prog Biophys Mol Biol 2021;162:101–10.

[28] Fedosejevs CS, Schneider MF. Sharp, localized phase transitions in single neuronal cells. Proc Natl Acad Sci 2022;119:e2117521119.

[29] Klymchenko AS. Fluorescent probes for lipid membranes: From the cell surface to organelles. Acc Chem Res 2022;56:1–12.

[30] Lippert E von. Spektroskopische Bestimmung des Dipolmomentes aromatischer Verbindungen im ersten angeregten Singulettzustand. Zeitschrift Für Elektrochemie, Berichte Der Bunsengesellschaft Für Phys Chemie 1957;61:962–75.

[31] Weber G, Farris FJ. Synthesis and spectral properties of a hydrophobic fluorescent probe: 6-propionyl-2-(dimethylamino) naphthalene. Biochemistry 1979;18:3075–8.

[32] Jin L, Millard AC, Wuskell JP, Dong X, Wu D, Clark HA, et al. Characterization and Application of a New Optical Probe for Membrane Lipid Domains. Biophys J 2006;90:2563–75. 10.1529/biophysj.105.072884.

[33] Danylchuk DI, Moon S, Xu K, Klymchenko AS. Switchable solvatochromic probes for live-cell super-resolution imaging of plasma membrane organization. Angew Chemie 2019;131:15062–6.

[34] Klymchenko AS. Solvatochromic and fluorogenic dyes as environment-sensitive probes: design and biological applications. Acc Chem Res 2017;50:366–75.

[35] Danylchuk DI, Sezgin E, Chabert P, Klymchenko AS. Redesigning solvatochromic probe laurdan for imaging lipid order selectively in cell plasma membranes. Anal Chem 2020;92:14798–805.

[36] Parasassi T, Conti F, Gratton E. Time-resolved fluorescence emission spectra of Laurdan in phospholipid vesicles by multifrequency phase and modulation fluorometry. Cell Mol Biol 1986;32:103–8.

[37] Parasassi, T and De Stasio, G and Ravagnan, G and Rusch, RM and Gratton E. Quantitation of lipid phases in phospholipid vesicles by the generalized polarization of Laurdan fluorescence. Biophys J 1991;60:179–89.

[38] Owen DM, Rentero C, Magenau A, Abu-Siniyeh A, Gaus K. Quantitative imaging of membrane lipid order in cells and organisms. Nat Protoc 2012;7:24–35.

[39] Mazeres S, Joly E, Lopez A, Tardin C. Characterization of M-laurdan, a versatile probe to explore order in lipid membranes. F1000Research 2014;3.

[40] Heimburg T. Thermal biophysics of membranes. John Wiley & Sons; 2008.

[41] Chong PLG. Effects of hydrostatic pressure on the location of PRODAN in lipid bilayers and cellular membranes. Biochemistry 1988;27:399–404. 10.1021/bi00401a060.

[42] Bacalum M, Radu M, Osella S, Knippenberg S, Ameloot M. Generalized polarization and time-resolved fluorescence provide evidence for different populations of Laurdan in lipid vesicles. J Photochem Photobiol B Biol 2024;250:112833. 10.1016/j.jphotobiol.2023.112833.

[43] Mamdouh Z, Giocondi M-C, Le Grimellec C. In situ determination of intracellular membrane physical state heterogeneity in renal epithelial cells using fluorescence ratio microscopy. Eur Biophys J 1998;27:341–51.

[44] Niko Y, Didier P, Mely Y, Konishi G, Klymchenko AS. Bright and photostable push-pull pyrene dye visualizes lipid order variation between plasma and intracellular membranes. Sci Rep 2016;6:18870.

[45] Lee AG. Lipid–protein interactions in biological membranes: a structural perspective. Biochim Biophys Acta - Biomembr 2003;1612:1–40. 10.1016/S0005-2736(03)00056-7.

[46] Lepock JR. Measurement of protein stability and protein denaturation in cells using differential scanning calorimetry. Methods 2005;35:117–25. 10.1016/j.ymeth.2004.08.002.

[47] Silvius JR. Role of cholesterol in lipid raft formation: lessons from lipid model systems. Biochim Biophys Acta - Biomembr 2003;1610:174–83. 10.1016/S0005-2736(03)00016-6.

[48] Olsen BN, Bielska AA, Lee T, Daily MD, Covey DF, Schlesinger PH, et al. The structural basis of cholesterol accessibility in membranes. Biophys J 2013;105:1838–47.

[49] Mabrey S, Mateo PL, Sturtevant JM. High-sensitivity scanning calorimetric study of mixtures of cholesterol with dimyristoyl-and dipalmitoylphosphatidylcholines. Biochemistry 1978;17:2464–8.

[50] Demel RA, De Kruyff B. The function of sterols in membranes. Biochim Biophys Acta (BBA)-Reviews Biomembr 1976;457:109–32.

[51] Burns M, Wisser K, Wu J, Levental I, Veatch SL. Miscibility Transition Temperature Scales with Growth Temperature in a Zebrafish Cell Line. Biophys J 2017;113:1212–22. 10.1016/j.bpj.2017.04.052.

[52] Crowe JH, Hoekstra FA, Crowe LM, Anchordoguy TJ, Drobnis E. Lipid phase transitions measured in intact cells with fourier transform infrared spectroscopy. Cryobiology 1989;26:76–84. 10.1016/0011-2240(89)90035-7.

